# Partitioning protein ParP directly links chemotaxis to biofilm dispersal in *Pseudomonas aeruginosa*

**DOI:** 10.1101/330878

**Authors:** Jesse M. Reinhardt, Sonia L. Bardy

## Abstract

The recent characterization of partitioning proteins in the localization of chemotaxis signal transduction systems was proposed to have broad implications for polarly-flagellated non-enterobacteriaceae gamma-proteobacteria. These studies showed that the loss of either partitioning protein resulted in equivalent reductions in swimming motility and chemotaxis protein localization and inheritance. However, the role of these chemotaxis partitioning proteins outside of *Vibrio spp*. remains untested. Our studies on the chemotaxis partitioning proteins in *Pseudomonas aeruginosa* revealed an unexpected role for the partitioning protein ParP. While the *P. aeruginosa* ParC and ParP homologs are needed for wild type swimming motility, the loss of ParP results in a greater swimming defect compared to the *parC* mutant. Our studies revealed that the Par-like proteins directly interact with each other and the chemotaxis system, and ParP interacts with DipA. Deletion of *dipA* results in a similar defect in swimming motility as the *parP* mutant. ParP has an interdependence for polar cluster formation, but not localization, with both CheA and DipA, and CheA cluster formation is partially dependent on ParP. Due to the direct interactions and interdependence of cluster formation of ParP and DipA, and the similar phenotypes of the *parP* and *dipA* mutants, further investigation into the role of ParP in biofilm dispersion is warranted.

**Importance:** Impaired chemotaxis protein cluster formation or inheritance reduces chemotaxis which can have an impact on of the virulence of a bacterium. In some gamma-proteobacteria there are systems in place to ensure that chemotaxis proteins, like chromosomes and plasmids, are localized for optimal chemotaxis and that daughter cells inherit their own clusters for use after cell division. Par-like proteins have been implicated in the partitioning and localization of chemotaxis proteins and the chemotactic ability of *Vibrio spp*. and *Rhodobacter sphaeroides* [1–3]. We propose that Par-like proteins can do more than localize chemotaxis proteins to the poles of the cells. In *P. aeruginosa*, they bring together other key proteins involved in regulating flagellar-based motility, and we propose they function as a critical link between biofilm dispersal and motility.

## Introduction

Spatial organization within bacterial cells results in the arrangement of proteins in distinct subcellular locations. This organization is not always static, and in some instances, can change in response to external cues or different stages within the bacterial lifecycle [4, 5]. There are a significant number of cellular processes that are affected by spatial organization and polarity, including signal transduction and motility. Bacterial chemotaxis is mediated by a two-component chemosensory system wherein a motile bacterium senses chemo-effectors in its environment and responds by moving towards favorable or away from unfavorable conditions. This system is best-studied in *Escherichia coli* where upon ligand binding, transmembrane methyl-accepting chemotaxis proteins (MCPs) will transduce the signal across the cytoplasmic membrane to the chemotaxis histidine kinase, CheA. CheA and the scaffolding protein CheW interact with the signaling domain of the MCP. The activation of CheA results in trans-autophosphorylation and transfer of the phosphate group to the response regulators CheY or CheB [6]. Phosphorylated CheY diffuses to the flagellar motor to cause a change in flagellar rotation, which results in a random change in swimming direction [7].

Specific localization patterns are known to be critical for optimal signal transduction [1, 8]. In *Vibrio cholera*, a polarly-flagellated gamma-proteobacterium, polar chemotaxis protein clusters are required for chemotaxis [1]. The Par-like proteins are required for proper cluster formation and localization of the polar chemotaxis proteins [1, 2]. ParC and ParP are homologous to ParA and ParB, which are partitioning (Par) proteins that are used for partitioning plasmids and chromosomes upon cell division. Deletion of the Par-like proteins of *V. cholerae* altered flagellar rotation, swimming motility, and chemotaxis protein localization [1, 2]. Specifically, deletion of *parC* or *parP* from *Vibrio parahaemolyticus* resulted in a ∼25-30% decrease in swimming motility and ∼50-60% of cells either having aberrant chemotaxis protein localization or partitioning [2]. The Par proteins mark the old pole and are recruited through the diffusion and capture of ParC by a HubP-dependent anchor. ParP interacts with both MCPs and CheA, thereby stimulating array formation [9]. These chemotaxis partitioning proteins are conserved in all polarly-flagellated gamma-proteobacteria [1].

*Pseudomonas aeruginosa*, another Gram-negative polarly-flagellated gamma-proteobacterium, is ubiquitous in the environment and commonly found in water, soil and on man-made structures [10]. It can act as an opportunistic pathogen and significantly contributes to morbidity and mortality in chronic infections in Cystic Fibrosis patients [11]. In *P. aeruginosa*, the chemotaxis system controlling swimming motility is encoded in two gene clusters, cluster I (*che I*) and cluster V, and the encoded proteins localize to the poles [12, 13]. Within *che I* are genes encoding for ParC and ParP homologs that may have importance in swimming motility and chemotaxis protein localization.

Spatial organization in *P. aeruginosa* is driven through a polar determinant called the polar organelle coordinator, or POC, complex for the flagellum, type IV pili, and chemotaxis proteins [14]. The POC complex consists of three proteins: TonB3, PocA, and PocB, which are currently known to sit at the top of the flagellar localization hierarchy above FlhF [14]. In *tonB3*, *pocA*, and *pocB* mutants, FlhF, CheA, and the flagellum are mislocalized from the cell pole. After the POC complex, FlhF is above all other known proteins for polar flagellar localization [15, 16]. Deletion of *flhF* results in mislocalized chemotaxis proteins and flagella, and reduced swimming motility [14]. Aside from FlhF and the Poc complex, there are no other major polar determinants of the chemotaxis system proteins known in *P. aeruginosa*.

Motility of *P. aeruginosa* is also affected by levels of bis-(3’→5’)-cyclic dimeric guanosine monophosphate (c-di-GMP), a bacterial second messenger that also regulates biofilm formation and dispersion, differentiation, and virulence [17, 18]. In regards to chemotaxis and biofilm formation, c-di-GMP levels are widely known to dictate the switch between motile (planktonic) and sessile (biofilm) states of growth. While there are obvious benefits to growing in a biofilm, bacterial cells can revert to planktonic growth. Environmental signals such as glutamate or succinate trigger *P. aeruginosa* to switch from a biofilm to a planktonic mode of growth – this process is known as biofilm dispersion. Biofilm dispersion requires DipA, a c-di-GMP phosphodiesterase, to mediate a cellular reduction of c-di-GMP levels [18–20].

DipA is also involved in chemotaxis and its absence results in defects in swimming and swarming motility in bulk population assays [21]. *P. aeruginosa* exhibits individual cell c-di-GMP heterogeneity due to the asymmetrical inheritance of DipA [16]. Most *dipA* mutant cells have high levels of c-di-GMP, which results in an overall reduction in average cell velocity and flagellar reversals. DipA is polarly-localized and forms a complex with the flagellum and CheA. The localization of DipA is completely dependent on the presence of the chemotaxis histidine kinase CheA and the phosphorylation of CheA promotes DipA PDE activity [16].

In our studies, we determined what effect the loss of the Par-like proteins has on swimming motility and chemotaxis protein cluster formation and localization in *P. aeruginosa* PAO1. We performed a bacterial two-hybrid assay to identify proteins that interact with the Par-like proteins. Finally, we examine the interdependence on cluster formation of the Par-like proteins with CheA and DipA. Our experiments suggest that the Par-like protein ParP is involved in the recruitment of the biofilm dispersal protein DipA to the flagellated pole.

## Results

### Par-like proteins are required for optimal chemotaxis in *Pseudomonas aeruginosa*

The chemotaxis gene cluster *che I* of *P. aeruginosa* encodes most of the genes required for chemotactic control of flagellar-based motility [13]. This includes the *par*-like genes *parC* and *parP* (Fig. 1A). Homologs of these genes are found in other polarly-flagellated non-Enterobacteriaceae γ-proteobacteria [2]. In *V. parahaemolyticus*, individual or double deletions of *parC*_*Vp*_ and *parP*_*Vp*_ resulted in a ∼25-30% defect in swimming motility. This swimming defect was due to an increase in the percentage of the cell population that lack chemotaxis protein foci or have mislocalized (non-polar) chemotaxis protein foci [2]. Due to the amino acid sequence homology between ParC and ParP in *V. parahaemolyticus* and *P. aeruginosa* and the conserved genetic organization surrounding these genes, ParC and ParP were proposed to be important for swimming motility in *P. aeruginosa*. Deletion of *parC* and *parP* resulted in a 25% and 70% reduction in swimming motility, respectively, and could be partially complemented through expression of His-tagged fusion proteins (Fig. 1B). These results suggested that ParP has a more important role in chemotaxis than ParC. Negative controls for swimming motility and chemotaxis are provided by the *fliC::tn* mutant and cluster I mutant respectively. The *par*C and *parP* mutants have a similar growth rate as wild type, demonstrating that the swimming defect is not due to a growth defect (data not shown). Given that the swimming defect seen in the *parP*_*Pa*_ mutants was significantly different from that reported in *Vibrio* [2], we further investigated the roles of these partitioning proteins in *P. aeruginosa*.

**Fig. 1.**
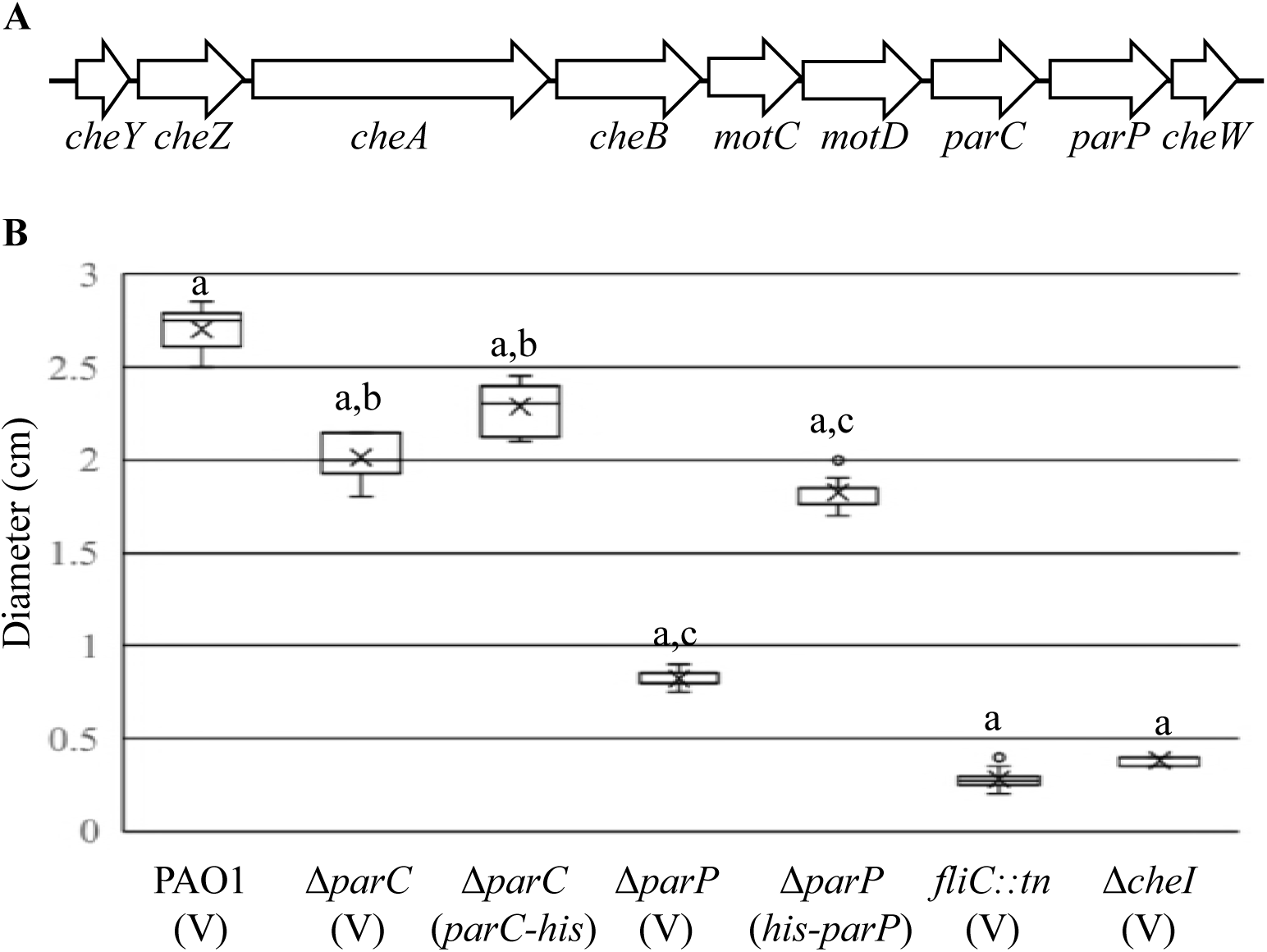
Par-like proteins encoded within chemotaxis gene cluster I (*che I*) are required for optimal swimming motility. A) *che I* gene cluster of *P. aeruginosa* (drawn to scale). B) Swimming motility assay of wild type (PAO1) and indicated *P. aeruginosa* strains. Δ*che I* strain lacks the chemotaxis genes *cheY*, *cheZ*, *cheA*, *cheB*, and *cheW*. The average swimming diameter measurements are shown and error bars denote the standard error of the mean. Matching letters indicate statistically significant differences, p<0.001. (V) indicates empty vector (pJN105).

### Chemotaxis protein localization is dependent on the Par-like proteins

To determine the cause of the swimming motility defects in the *parC*_*Pa*_ and *parP*_*Pa*_ mutants, we examined CheA (histidine kinase) localization and expression. The chemotaxis proteins of *P. aeruginosa* normally localize to the poles of the cell [22]. It has been previously demonstrated in *V. parahaemolyticus* that deletion of *parC*_*Vp*_, *parP*_*Vp*_, or both resulted in 50-60% of cells having a reduction in either chemotaxis protein foci formation or polar localization [2]. Through fluorescence microscopy, it was determined that in *P. aeruginosa* ParC and ParP were required for wild type levels of chemotaxis protein foci formation (Fig. 2). CheA-mTurquiose (CheA-mTq) expressed from the native site in the chromosome was used as a marker for chemotaxis protein foci formation and localization [16] as CheA, along with CheW and MCPs, are required for higher order clustering [23]. As a control, CheA foci formation was tested in the *cheW* mutant and showed a 96% reduction as previously published [22]. CheA foci formation was reduced by ∼45-50% in the *parC*_*Pa*_ and *parP*_*Pa*_ deletion mutants (Fig. 2B). Surprisingly, in the *parC*_*Pa*_ and *parP*_*Pa*_ deletion strains, the localization of CheA foci remained largely unchanged compared to wild type. This suggests that the Par-like proteins are more important for foci stability or inheritance as opposed to localization in *P. aeruginosa*. The three amino acid residues that are different between CheA from PAO1 and PA14 do not affect function as the *P. aeruginosa* PAO1 strain expressing CheA-mTq from PA14 was capable of wild type chemotaxis (Fig. 2D) and therefore its use was justified for localization studies. The CheA-mTq fusion protein was present in all mutant backgrounds (Fig. 2B), demonstrating that the lack of foci formation was not due to reduced levels of CheA. Curiously, western blotting suggested that CheA-mTq levels were slightly higher in the mutants compared to wild type (Fig. 2C). The reason for this increase in CheA levels remains to be determined.

**Fig. 2.**
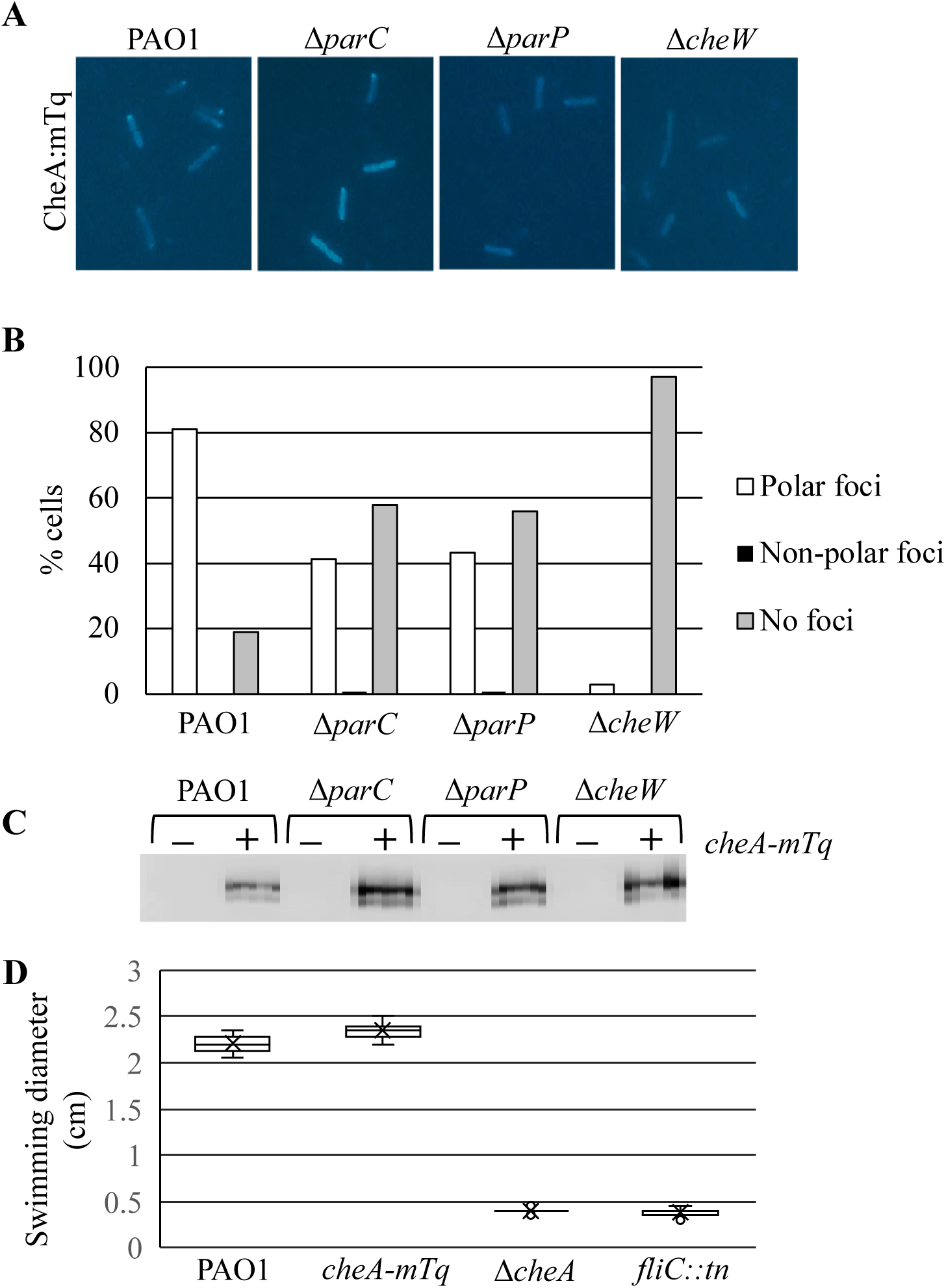
The Par-like proteins affect chemotaxis protein localization. A) Representative images of CheA:mTq foci formation in wild type (PAO1) and indicated *P. aeruginosa* strains. B) Quantitation of CheA:mTq foci localization in the indicated *P. aeruginosa* strains. 248 cells per strain were counted. C) CheA:mTq expression levels as determined through western blotting. D) Expression of CheA:mTq supports swimming motility. Average swimming diameter is shown and error bars denote the standard error of the mean. All values were significantly different from wild type, p<0.001.

### DipA interacts with ParP_Pa_ and affects swimming motility

Because deletion of the *par*-like genes affected swimming motility and chemotaxis protein foci formation in our work and in the recent studies in *Vibrio spp*., we investigated protein interactions between ParC_Pa_ and ParP_Pa_ as well as chemotaxis proteins and MCPs [1, 2, 9]. Given that the genome of *P. aeruginosa* is reported to encode 23 MCPs that feed into the flagellar based chemotaxis system, a single representative MCP (PA2867) was assayed for interaction with the Par-like proteins. A bacterial two-hybrid (B2H) assay showed that ParC_Pa_ and ParP_Pa_ directly and strongly interact with each other and weakly interact with CheA and the MCP (Fig. 3). ParC_Pa_ could self-interact, thus further suggesting that it is acting as a ParA-like protein [2, 24].

**Fig. 3.**
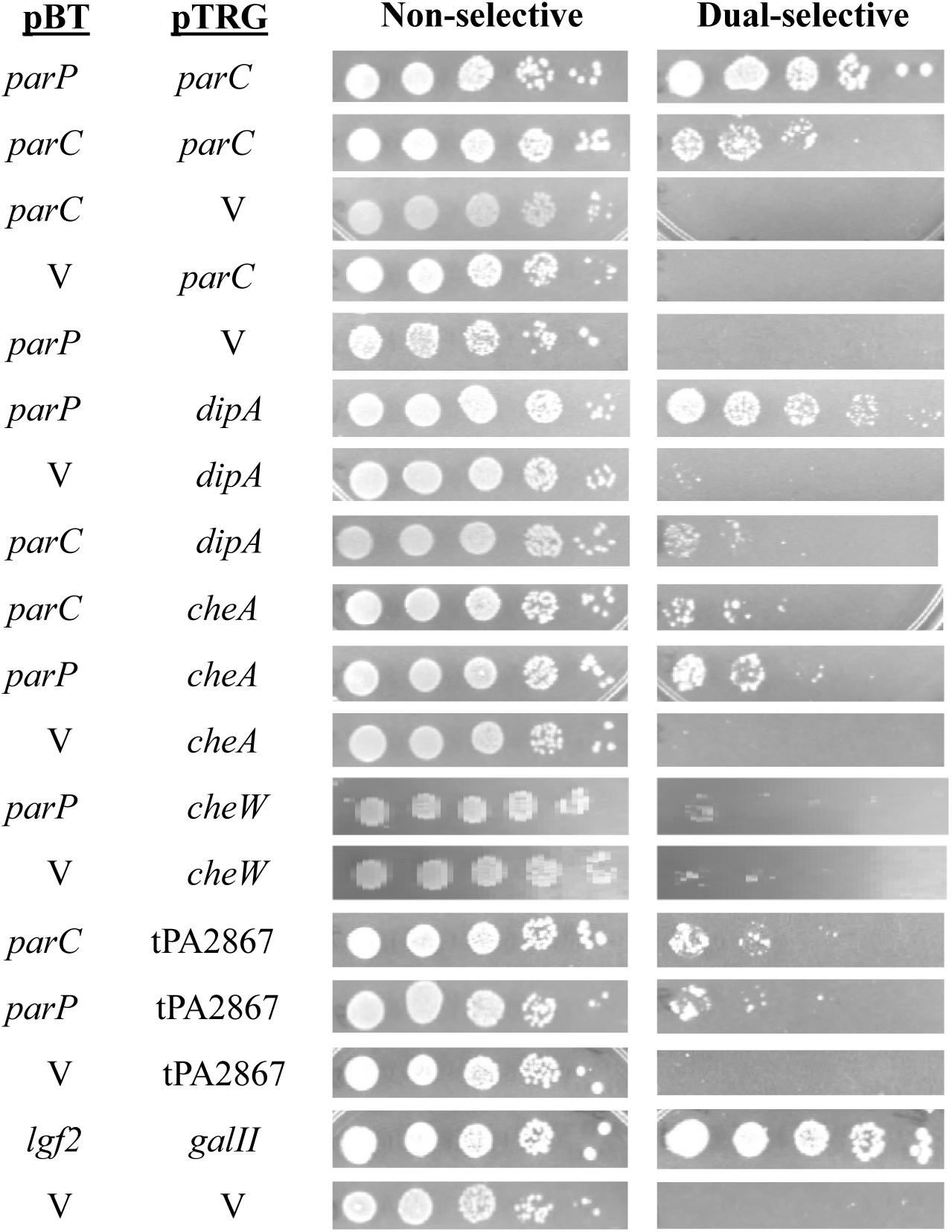
DipA interacts directly with ParP, as demonstrated by a bacterial two-hybrid assay. 5 µl of a 10-fold dilution series are spotted from left to right. Cultures on the non-selective media function as a loading control, while dual-selective media reveals the strength of the protein-protein interactions. Strong interactions have growth to the right-most spot, as indicated by the positive control *lgf2* and *galII*.

It was previously reported in *P. aeruginosa* strain PA14, that CheA co-immunoprecipitated with the phosphodiesterase PA5017 (hereafter referred to as DipA for clarity within the literature) [16]. This indicated that CheA and DipA form a complex with each other, but it was not known if this interaction was direct or indirect. DipA is known to be involved in biofilm dispersion and swimming motility and its ability to form polar foci and degrade c-di-GMP is dependent on CheA [16, 18, 20]. Because the Par-like proteins affect swimming motility and CheA foci formation, DipA and the Par proteins were assayed for direct interactions to determine if the Par proteins aid in CheA/DipA complex formation. Strikingly, a B2H assay revealed that ParP_Pa_ directly and strongly interacts with DipA (Fig. 3). No direct interaction could be detected between DipA and CheA or ParP and CheW using this assay.

In agreement with previous studies, the *dipA* mutant showed a 63% reduction in swimming motility [16]. This was similar to the 70% reduction in swimming motility seen in the *parP*_*Pa*_ mutant, yet these results were significantly different from each other (Fig. 4A). Complementation with His-DipA fully restored swimming motility to the *dipA* mutant (Fig. 4A). A Western blot confirmed that His-DipA was expressed (Fig. 4B).

**Fig. 4.**
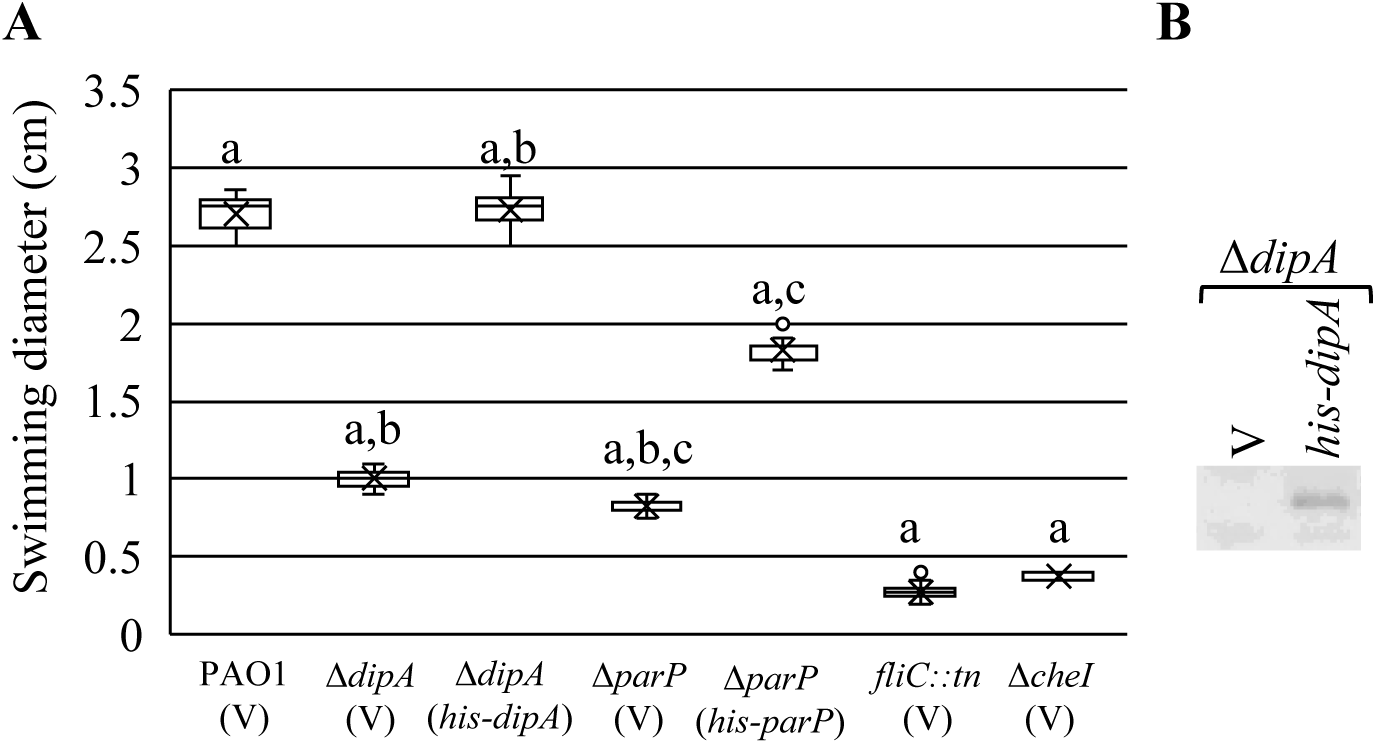
Deletion of DipA results in a similar reduction of swimming motility as seen in Δ*parP*. (A) Swimming motility assay of indicated *P. aeruginosa* strains. The averaged swimming diameters are shown and error bars denote standard error of the mean. Matching letters indicate statistically significant differences, p>0.001. (B)Western blot showing expression of His-DipA.

### DipA, ParP_Pa_ and CheA polar localization is interdependent

Given the similar phenotypes and direct interaction between ParP and DipA, we investigated the localization dependence of CheA, DipA and ParP_Pa_ by fluorescence microscopy. CheA-mTq foci formation or localization remained unchanged in a *dipA* mutant, indicating that CheA localization is independent of DipA (Figs. 5A and B). Levels of CheA-mTq remained unchanged in the *dipA* mutant (Fig. 5C). Yfp-ParP_Pa_ foci formation was reduced by 50% in a *dipA* mutant and 60% in a *cheA* mutant, but there was no change in localization (Fig. 6). DipA-Yfp foci formation was reduced by 50% in a *parP*_*Pa*_ mutant and 95% in a *cheA* mutant (Fig. 7). The dependence of DipA on CheA for foci formation has been previously published [16]. Expression of the ParP and DipA fluorescent fusion proteins complemented the swimming defect of their respective mutant parent strains to the same levels as the His-tagged ParP and DipA proteins (data not shown), thereby demonstrating that these fusion proteins are as functional as the His-tagged versions. DipA was present at similar levels in all mutant backgrounds, demonstrating that a loss of foci formation was not due to altered protein levels (Fig. 7B). The levels of ParP fusion protein in the Δ*parP*Δ*cheA* and Δ*parP*Δ*dipA* double deletion strains consistently appeared less than in Δ*parP*, suggesting ParP stability may be affected by the loss of its interacting partners (Fig. 6B). Combined, the results of these fluorescence microscopy studies on ParC, ParP, CheA and DipA localization show that there is an interdependence on localization, particularly for ParP on CheA and DipA, DipA on ParP, and CheA on ParP (Fig. 8).

**Fig. 5.**
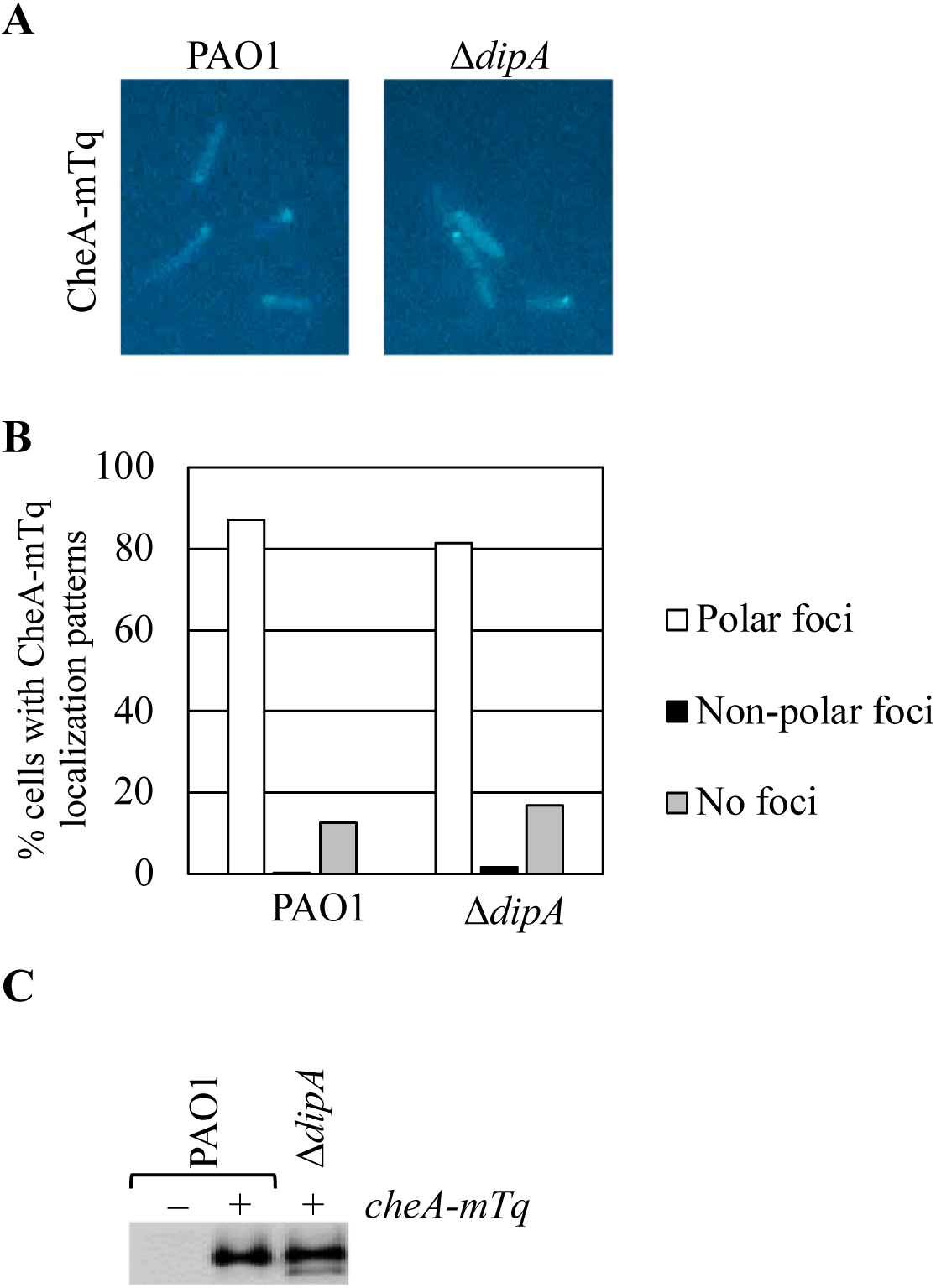
DipA is not required for CheA foci formation or localization. (A) Representative images of CheA-mTq foci formation in wild type and mutant *P. aeruginosa* strains. (B) Quantitation of CheA-mTq foci formation and localization in the indicated *P. aeruginosa* strains. 300 cells were counted per strain. (C) Western blot showing CheA-mTq levels.

**Fig. 6.**
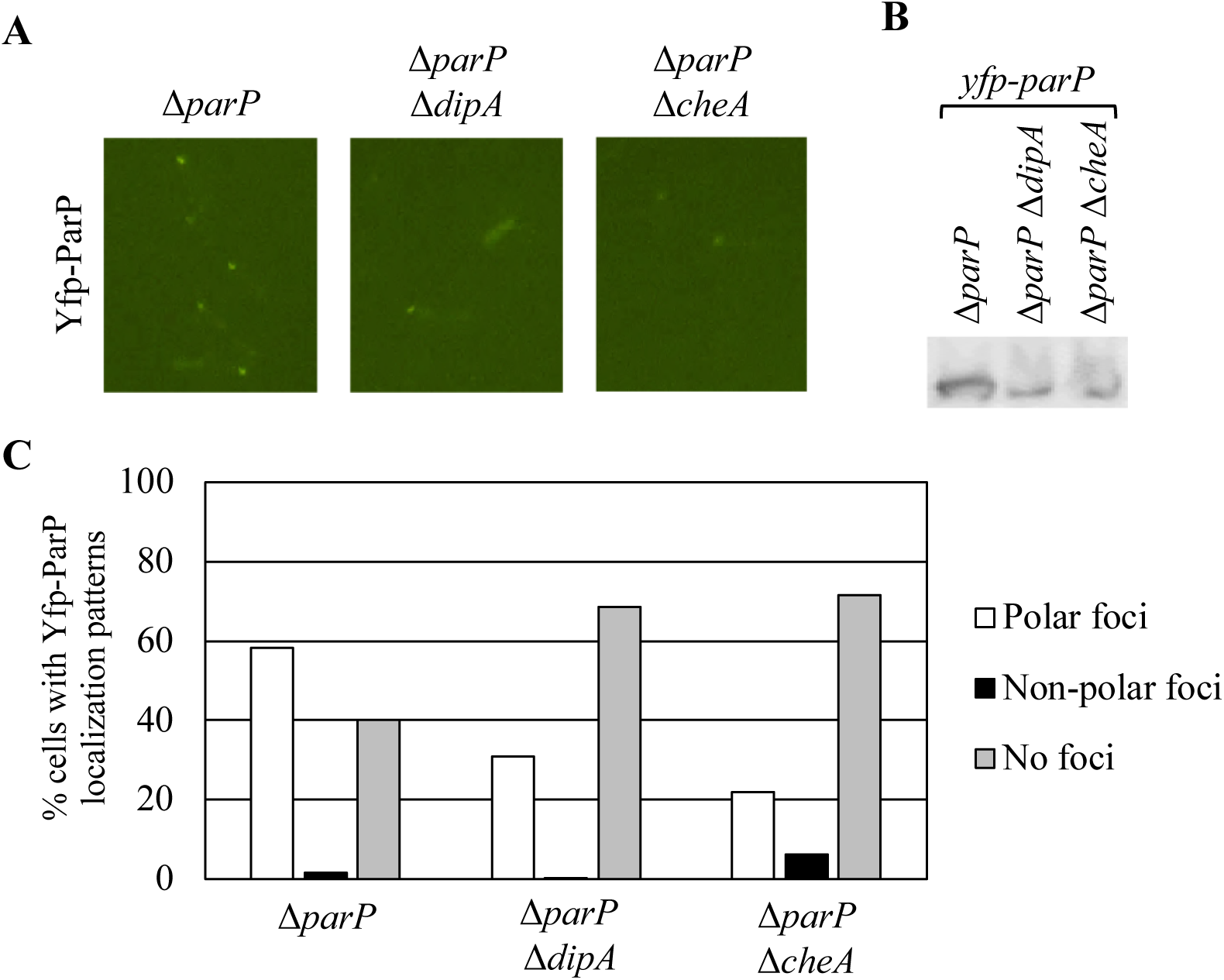
DipA and CheA influence ParP foci formation. (A) Representative images of Yfp-ParP foci formation in wild type and mutant *P. aeruginosa* strains. (B) Western blot showing Yfp-ParP levels in the indicated strains. (C) Quantitation of Yfp-ParP foci formation and localization patterns in the indicated *P. aeruginosa* strains. 300 cells were counted per strain.

**Fig. 7.**
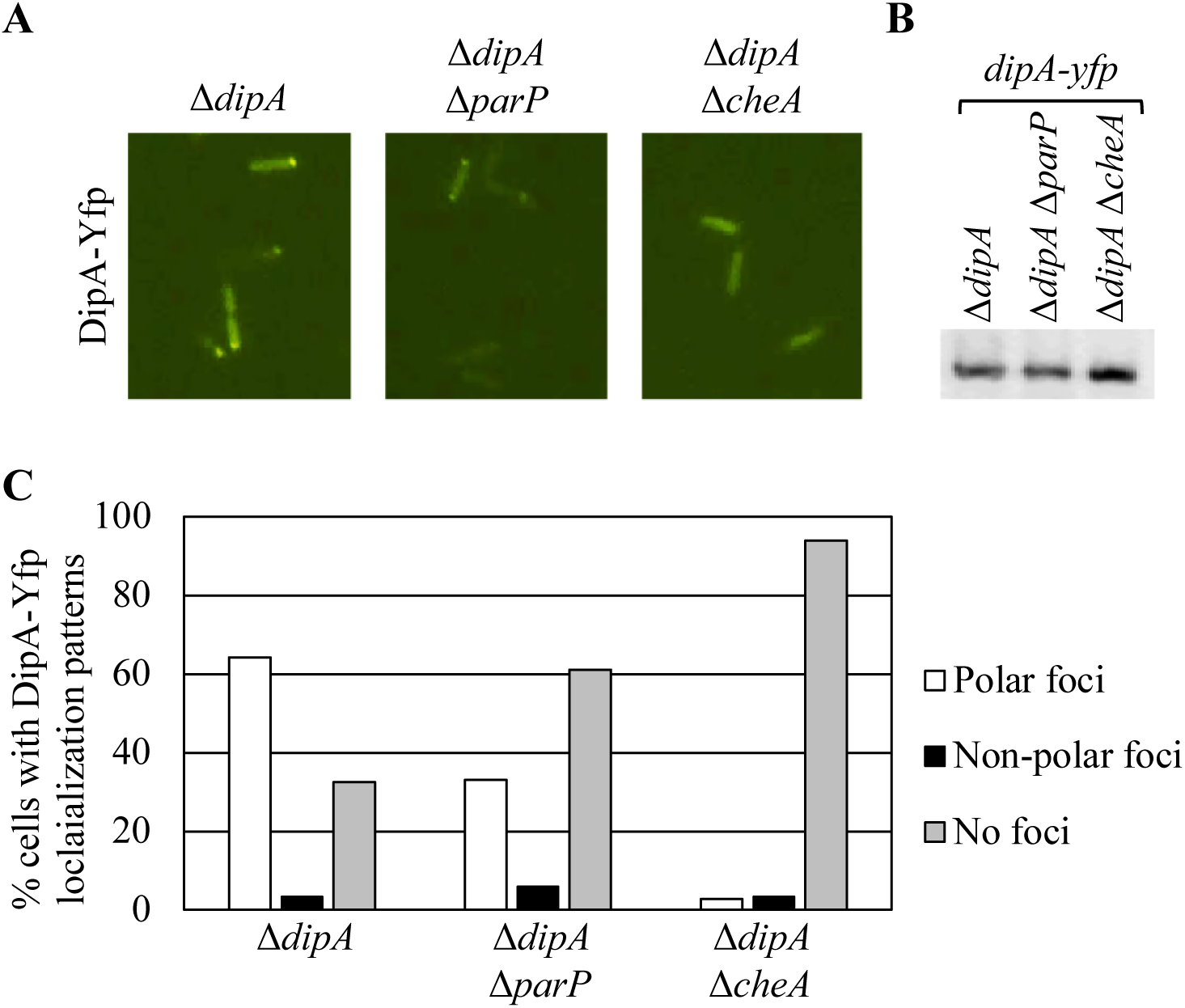
DipA foci formation is influenced by ParP and dependent on CheA. (A) Representative images of DipA-Yfp foci formation in wild type and mutant *P. aeruginosa* strains. (B) Western blot showing DipA-Yfp levels in the indicated strains. (C) DipA-Yfp foci formation and localization patterns in the indicated *P. aeruginosa* strains. 300 cells were counted per strain.

**Fig. 8.**
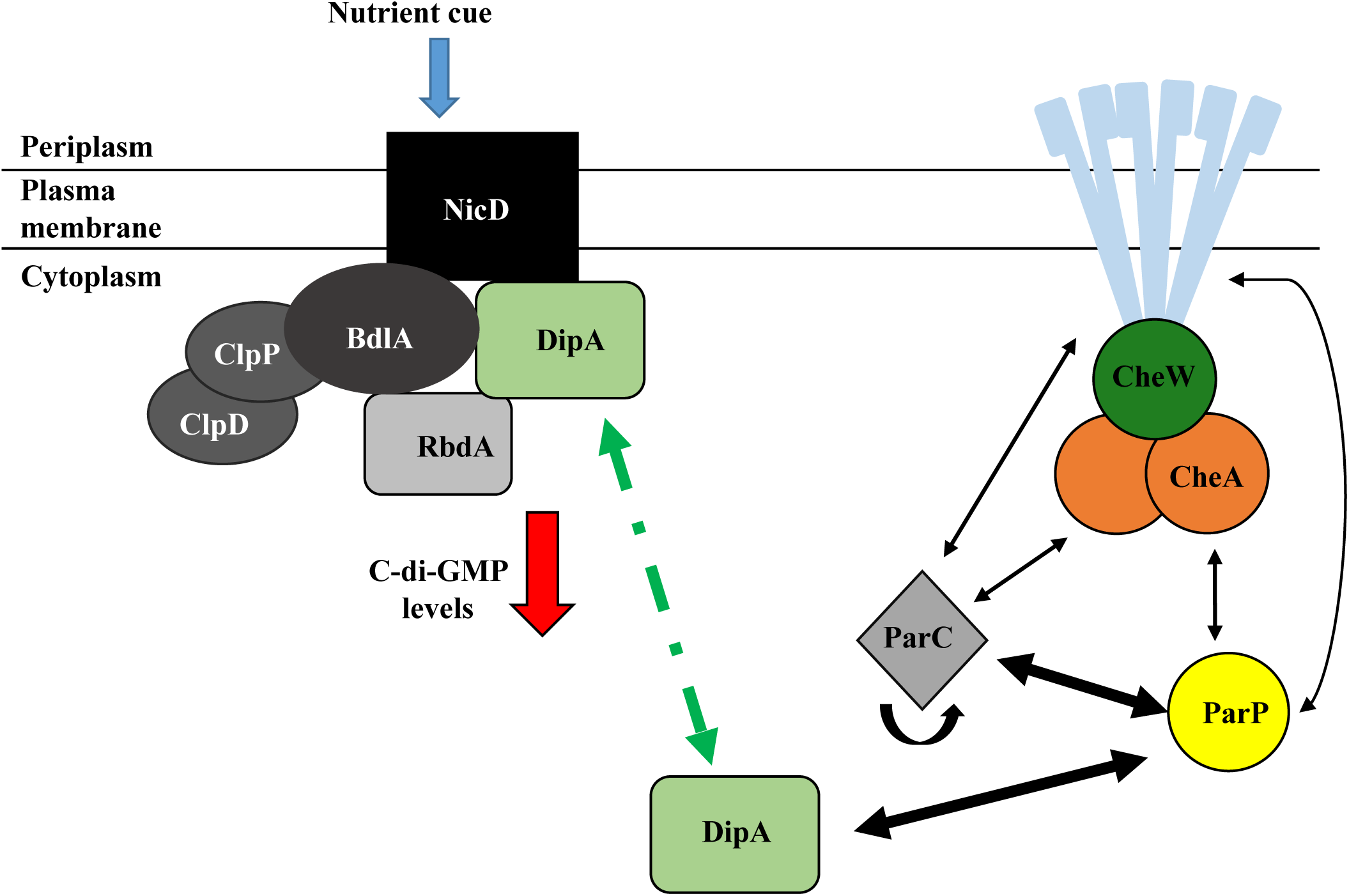
Model showing B2H interactions linking the Par-like proteins with the chemotaxis and biofilm dispersion systems of *P. aeruginosa*. Black arrows indicate direct protein-protein interactions, with thicker arrows being a stronger interaction. The green dashed arrow points to the different roles that DipA has in regards to biofilm dispersion and chemotaxis. The red arrow pointing down indicates a decrease in c-di-GMP levels. The blue arrow represents a nutrient cue that is sensed by NicD.

## Discussion

Chemotaxis proteins localize to distinct regions within a bacterial cell – this localization can vary depending on if it is a random or ordered process. A variety of mechanisms have been proposed including membrane curvature, stochastic nucleation, nuclear exclusion and interaction with the Tol-Pal complex [25–29]. These different mechanisms may not be exclusive of each other as these studies focus on different bacteria such as *E. coli* and *Bacillus subtilis*, as well as different MCPs within the same species. In *E. coli*, MCPs localize to the poles as large clusters, yet small clusters and individual proteins can be seen at the lateral regions of the inner membrane [26]. Other organisms, such as *Vibrio spp*. and *R. sphaeroides*, have *par*-like genes in their chemotaxis gene clusters and the encoded proteins are used for chemotaxis protein cluster formation and localization [1–3]. *P. aeruginosa* has *par*-like genes encoded in its main chemotaxis gene cluster, *che I* (Fig. 1A), and this work provides convincing evidence that these Par-like proteins are involved in chemotaxis and linked to DipA, a c-di-GMP phosphodiesterase involved in biofilm dispersion and motility regulation.

The Par-like proteins are known to be involved in swimming motility, chemotaxis and polar array formation in *Vibrio* spp. [1, 2, 9]. Our work shows that in *P. aeruginosa*, ParC_Pa_ and ParP_Pa_ are needed for optimal swimming motility (Fig. 1B). Comparison of the phenotypes between *V. parahaemolyticus* and *P. aeruginosa* reveal that the *parP*_*Vp*_ mutant has a swimming defect equal to the *V. parahaemolyticus parC* mutant. However, ParP_Pa_ is distinct in that it appears to have a more significant role in swimming motility, and possible reasons for this will be discussed below.

The Par-like proteins are known to dimerize and interact with each other and with the chemotaxis system via CheA and the MCPs in *V. parahaemolyticus* [2, 9]. Our work confirms that ParC_Pa_ can dimerize and strongly interact with ParP_Pa_, and both proteins interact with CheA (Fig. 3). We did not observe ParP_Pa_ self-interaction (data not shown). In agreement with earlier studies, it was determined that the Par-like proteins of *P. aeruginosa* interacted with a representative MCP, thus demonstrating that ParC_Pa_ and ParP_Pa_ are not linked to the chemotaxis system only via CheA [9]. Strikingly, we found that ParP_Pa_ interacted strongly with DipA (Fig. 3). These results are novel, as ParP_Pa_ and DipA form the first direct link between the biofilm dispersion and chemotaxis systems. It was previously shown by co-immunoprecipitation that DipA and CheA form a complex, but it was not known if this was through direct or indirect interactions [16]. Additionally, this earlier publication focused on the role of DipA (referred to as Pch) in motility and did not address the role of this phosphodiesterase in biofilm dispersal [16, 18].

The *dipA* mutant had a reduction in swimming motility that was similar, but significantly different to what was seen in the *parP*_*Pa*_ mutant (Fig. 4A). Localization studies suggest that the motility defect may be due to the loss of ParP as well as altered c-di-GMP levels at the cell pole. In *P. aeruginosa* PA14, Kulasekara *et al* [16] showed that loss of DipA leads to a loss of c-di-GMP heterogeneity in individual cells, with most cells having high levels of c-di-GMP. A reduction in flagellar reversals and average cell velocity compared with wild type was also observed. These results suggest that c-di-GMP levels modulate flagellar reversals and cell velocity, however, the mechanism by which this occurs has not been determined but may involve a c-di-GMP effector protein. DipA forms polar foci at the flagellated pole with CheA. After cell division, one of the daughter cells will inherit the flagellum and a DipA cluster, which lowers the c-di-GMP levels in that cell, thus creating c-di-GMP heterogeneity in individual cells. The role of this heterogeneity is speculated to give a survival advantage to these cells in unpredictable environments [16]. Individual cells with high or low c-di-GMP levels would likely tend to either attach to a surface and start biofilm formation or remain motile and spread to new areas. In this sense, at any moment, there are cells that are “primed” for either choice, depending on the environment. The presence of CheA is absolutely required for DipA polar localization and the phosphorylation activity of CheA promotes DipA PDE activity. The GTPase FlhF is required for polar localization of the flagellum, and in an *flhF* mutant, the flagellum is still produced but mislocalized from the pole [30]. This results in cells having reduced swimming and swarming motility. Loss of FlhF also results in a reduction of transcription of class II, III or IV flagellar genes [30]. FlhF is above CheA and DipA in terms of dictating polar localization, but not their association with each other [16]. The absence of FlhF results in the mislocalization of the flagellum, and CheA and DipA foci from the pole. This suggests that the flagellum, CheA and DipA form a complex at one pole of the cell. However, it is not known if these three components remain in a complex when they are mislocalized from the pole. By forming these protein complexes, new daughter cells will be more likely to inherit necessary chemotaxis proteins to be used right away or once they synthesize a new flagellum.

Using fluorescence microscopy, we tested chemotaxis protein localization in the absence of the Par-like proteins. Deletion of either ParC_Pa_, ParP_Pa_ or CheW resulted in a loss of CheA cluster, or foci, formation, but not polar localization in *P. aeruginosa* (Fig. 2). Comparable results were seen for the Par-like proteins in *V. parahaemolyticus*, except that of the cells that had aberrant clustering, 50% of them had no clusters while the other 50% had non-polar clusters [2]. These results suggest that in *P. aeruginosa*, the Par-like proteins function more in cluster stability as opposed to localization, but we cannot rule out technical differences as the cause of this discrepancy. Our results for the loss of CheA cluster formation in a *cheW* mutant agree with previously published work [22]. Interestingly, we show that the loss in CheA cluster formation also coincided with a slight increase in CheA levels in the cells (Fig. 2C). The absolute levels of MCP, CheW and CheA proteins can vary in a bacterium, but their stoichiometry appears to remain constant [6, 31]. Overexpression of a chemotaxis protein can reduce chemotaxis and cluster formation [31, 32]. One possible explanation for the reduction in CheA cluster formation in *P. aeruginosa* is that excess levels of CheA are present in the cell relative to the MCP and CheW proteins. However, our results do not show if the stoichiometry of MCP:CheW:CheA was altered - this would require further investigation.

The Par-like proteins are interdependent in their polar cluster formation. ParC_Vp_ and ParP_Vp_ are both needed for their cluster formation and polar localization in *V. parahaemolyticus* [2]. While we have not tested the interdependence of ParC_Pa_ and ParP_Pa_, our work has shown that the clustering ability of ParP_Pa_ is interdependent on both CheA and DipA and that loss of cluster formation is ∼50% (Figs. 6 and 7). These results suggest that the interdependence of localization between these proteins are equally important in their cluster formation. In a previous study and in this work, DipA cluster formation requires CheA [16]. However, we found that CheA cluster formation and cellular levels are not dependent on DipA (Fig. 5).

In summary, this work showed that the Par-like proteins of *P. aeruginosa* PAO1 are involved in chemotaxis controlling swimming motility. Our results correlate well with other studies in terms of the effects of the Par-like proteins on swimming motility and chemotaxis protein foci formation. Notably, we found that ParP_Pa_ plays a more significant role in swimming motility than ParC_Pa_. We discovered that the c-di-GMP phosphodiesterase DipA directly interacts with ParP_Pa_ and that they have an interdependence in their cluster formation. These results suggest that ParP_Pa_ and DipA work in the same pathway and this may be the mechanism behind the large decrease in swimming motility in a *parP* mutant. We have provided compelling evidence that the chemotaxis and biofilm dispersion systems are linked together via DipA and ParP_Pa_ (Fig. 8). When biofilm cells sense a nutrient cue to disperse, *dipA*, motility, and chemotaxis genes are upregulated, c-di-GMP levels decrease, the extracellular matrix is broken down, and cell adhesiveness is reduced [18, 20]. Due to this series of events, cells become motile and chemotactic, and leave the biofilm. This leads to the question of what role ParP_Pa_ has in this process of dispersion and if DipA proteins can temporally, and perhaps spatially, switch between interactions with biofilm dispersal proteins and chemotaxis proteins, or if there are functionally separate pools of this protein within the cell. To our knowledge, the localization of DipA has not yet been determined in biofilm or biofilm-dispersed cells. Future studies will determine in more detail how the loss of ParP_Pa_ has a greater defect in swimming motility than the loss of ParC_Pa_ and if ParP_Pa_ has a role in biofilm dispersion.

## Materials and Methods

### Strains, plasmids, growth conditions and media used

Lists of plasmids and strains used in this publication are in Supplemental Tables 1 and 2, respectively. All *P. aeruginosa* strains generated in the work are derived from *P. aeruginosa* PAO1 (Iglewski strain). Both *E. coli* and *P. aeruginosa* were grown in Lysogeny Broth (LB) with aeration and on LB 1.5% agar plates at 37°C. Antibiotics were used at the following concentrations as appropriate: 50 µg/mL of gentamycin and 70 µg/mL of tetracycline for *P. aeruginosa* and 15 µg/mL of gentamycin, 30 µg/mL of kanamycin, 25 µg/mL of chloramphenicol and 10 µg/mL of tetracycline for *E. coli*.

### Generation of deletion mutants and expression strains

In-frame gene deletions of *cheA* and *parC* were generated by homologous recombination using the suicide vectors pEX18Tc or pEX19Gm as previously described [33, 34]. Briefly, 1 Kb DNA fragments upstream and downstream of the genes of interest were PCR amplified and fused together through splicing by overlap extension (SOE) PCR using PAO1 DNA as template [35]. Primers are listed in Supplemental Table 3. Fusion constructs were sequenced to ensure no undesired mutations were introduced. This resultant fragment was cloned into pEX18Tc or pEX19Gm and transformed into *E. coli* S17-1 for mating into *P. aeruginosa* PAO1. Merodiploids were selected on tetracycline or gentamycin, as appropriate, with chloramphenicol [5 µg/mL] providing counter-selection against *E. coli*. Resolution of the merodiploids was achieved through 10% sucrose counter-selection, and the deletions were confirmed by PCR. Gene deletions of *cheW*, *dipA*, and *parP* were performed as above except the upstream and downstream 1 Kb DNA fragments included nine basepairs from the 5’ and 3’ ends of the target gene. This approach was used to reduce the likelihood of polar effects.

Incorporation of *cheA-mTq* into the native site of the chromosome was done using a *cheA-mTq*:pEX19Gm construct [16] as above. In this construct, *cheA* from *P. aeruginosa* PA14 was used. The CheA amino acid sequences from strains PAO1 and PA14 are 99.6% identical, with three residues [E133A, A161V and P191S, respectively] being different between them. The *dipA-yfp* fusion was amplified, sequenced, and cloned into pJN105 [16, 36]. The *yfp-parP* fusion was generated by SOE PCR, sequenced and cloned into pJN105.

### Bacterial two-hybrid analysis

Protein interactions were tested using the BacterioMatch II Two-Hybrid System (Agilent Technologies). Briefly, the overnight cultures of the test strains were diluted to equal cell density. Five ten-fold serial dilutions of each culture were made and 5 µl of each was spotted on non-selective and dual-selective plates containing antibiotics and IPTG. The dual-selective plates had 5 mM 3-AT and 10 µg/ml streptomycin to test the strength of the protein interactions. The negative control strain harbored empty pBT and pTRG vectors, while the positive control strain harbored *lgf2*:pBT and *galII*:pTRG as supplied by the manufacturer. The pBT and pTRG constructs were made using standard cloning techniques. The pairwise interactions tested included ParP-ParP, ParC-ParC, ParP-DipA, ParC-DipA, ParP-CheA, ParP-CheW, ParC-PA2867_161-490_, and ParP-PA2867_161-490_. PA2867 is a transmembrane receptor/MCP, and so a truncated version (PA2867_161-490_ or tPA2867) containing only the C-terminal cytoplasmic portion was used.

### SDS-PAGE and western blot

Whole cell lysates were prepared from mid-late log (OD_600_ 0.5-1) liquid cultures (37°C, with aeration). The cells were harvested and suspended in 2X SDS loading buffer, and loading was normalized based on OD_600_. Whole cell lysates were separated by SDS-PAGE, and stained using Coomassie brilliant blue G-250 - perchloric acid solution [37]. The primary antibodies were α-His (1:3000), α-mCherry (1:1000) and α-GFP (1:1000). Secondary antibodies (1:10,000) conjugated to peroxidase allowed detection of signal using the SuperSignal West Femto Maximum Sensitivity Substrate kit. Western blots were visualized and imaged using a Fotodyne FOTO/Analyst FX system.

### Swimming assay

Fresh *P. aeruginosa* colonies were stab inoculated into swimming media (1% tryptone, 0.5% NaCl and 0.3% agar) with antibiotics as appropriate. Following incubation at 30°C for 18 hours, the diameter of the swimming zones were measured. For each assay, 12 biological replicates were performed. ANOVA calculations were followed by the Tukey HSD post-hoc test using the R Console program (Version 3.2.3).

### Fluorescence microscopy

Overnight cultures of *P. aeruginosa* were sub-cultured in LB broth with antibiotics as appropriate, and grown for three hours (with aeration at 37°C), resulting in cultures in mid-late log phase (OD_600_ 0.5 - 1). 5 µl of culture was spotted onto a polylysine-treated coverslip (Fisherbrand 25CIR-1D) for observation using a Nikon Eclipse 90*i* microscope with a Hamamatsu digital camera C11440 (ORCA-Flash 4.0) and a Nikon Intensilight C-GHFI halogen lamp. Images were captured under DIC, Yfp, and Cfp filters, as appropriate. For quantitation of localization patterns, between 248 and 300 cells were scored for foci formation and localization. Foci were labeled as being polar if they fell within the curvature of the poles or non-polar if they did not.

### Protein alignment

Clustal Omega multiple sequence alignment was used for comparing the amino acid sequences of multiple proteins [38].

## Acknowldegements

The authors thank Zachary Zawada and Ashton Novy for their contributions. We also thank Samuel Miller for indicated plasmid. The University of Washington *Pseudomonas aeruginosa* transposon mutant collection is supported by NIH P30 DK089507.

---

**Table S1:** Plasmids used in this study

| Plasmid | Description | Source |
| --- | --- | --- |
| $\Delta cheA$:pEX18Tc | DNA fusion product for deletion of *cheA* cloned into the EcoRI (5') and BamHI (3') sites of pEX18Tc | This study |
| $\Delta cheW$:pEX18Tc | DNA fusion product for deletion of *cheW* cloned into the EcoRI (5') and BamHI (3') sites of pEX18Tc | This study |
| $\Delta dipA$:pEX18Tc | DNA fusion product for deletion of *dipA* cloned into the EcoRI (5') and SacI (3') sites of pEX18Tc | This study |
| $\Delta parC$:pEX18Tc | DNA fusion product for deletion of *parC* cloned into the EcoRI (5') and BamHI (3') sites of pEX18Tc | This study |
| $\Delta parP$:pEX18Gm | DNA fusion product for deletion of *parP* cloned into the EcoRI (5') and HindIII (3') sites of pEX18Gm | This study |
| *cheA-mTq*:pEX18Gm | DNA fusion product for insertion of *cheA-mTurquoise* at the native chromosomal site, cloned into pEX18Gm | [16] |
| *dipA-yfp*:pUC18T-mini-TN7T-Gm | Plasmid template for amplifying *dipA-yfp* | [16] |
| pJN105 | Broad host range vector. pBBR-1 MCS5 AraC-pBAD derivative | [36] |
| his-*cheW*:pJN105 | his-*cheW* cloned into the EcoRI (5') and SacI (3') sites of pJN105 | This study |
| his-*dipA*:pJN105 | his-*dipA* cloned into the EcoRI (5') and XmaI (3') sites of pJN105 | This study |
| *parC*:pJN105 | *parC* cloned into the EcoRI (5') and XbaI (3') sites of pJN105 | This study |
| *parC*-his:pJN105 | *parC*-his cloned into the EcoRI (5') and XbaI (3') sites of pJN105 | This study |
| his-*parP*:pJN105 | his-*parP* cloned into the EcoRI (5') and SacI (3') sites of pJN105 | This study |
| *dipA-yfp*:pJN105 | *dipA-yfp* amplified from *dipA-yfp*:pUC18T-mini-TN7T-Gm and cloned into the EcoRI (5') and XbaI (3') sites of pJN105 | This study |
| *yfp-parP*:pJN105 | DNA fusion product *yfp-parP* cloned into the EcoRI (5') and SacI (3') sites of pJN105 | This study |
| pBT | Expression vector used for Bacterial Two-Hybrid | Agilent Technologies |
| pTRG | Expression vector used for Bacterial Two-Hybrid | Agilent Technologies |
| *cheA*:pTRG | *cheA* cloned into the BamHI (5') and EcoRI (3') sites of pTRG | This study |
| *dipA*:pBT | *dipA* cloned into the NotI (5') and EcoRI (3') sites of pBT | This study |
| *mcpS*:pTRG | *mcpS* cloned into the XhoI (5') and NotI (3') sites of pTRG | This study |
| *parC*:pBT | *parC* cloned into the NotI (5') and EcoRI (3') sites of pBT | This study |
| *parC*:pTRG | *parC* cloned into the NotI (5') and EcoRI (3') sites of pTRG | This study |
| *parP*:pBT | *parP* cloned into the NotI (5') and EcoRI (3') sites of pBT | This study |
| tPA2867:pTRG | tPA2867 cloned into the EcoRI (5') and XhoI (3') sites of pTRG | This study |

**Table S2:** Strains used in this study

| Strain | Description | Source |
| --- | --- | --- |
| *E. coli* BacterioMatch II Two-Hybrid System Reporter Strain | $\Delta(mcrA)183 \Delta(mcrCB-hsdSMR-mrr)173 endA1 hisB supE44 thi-1 recA1 gyrA96 relA1 lac [F' lacI^q HIS3 aadA Kan^r]$ | Agilent Technologies |
| *E. coli* XL-1 Blue MRF' kan<sup>r</sup> | $\Delta(mcrA)183 \Delta(mcrCB-hsdSMR-mrr)173 endA1 supE44 thi-1 recA1 gyrA96 relA1 lac [F' proAB lacI^q Z\Delta M15 Tn5 (Kan^r)]$ | Agilent Technologies |
| *E. coli* XL-1 Blue MRF' tet<sup>r</sup> | $recA1 endA1 gyrA96 thi-1 hsdR17 supE44 relA1 lac [F' proAB lacI^q Z\Delta M15 Tn10 (Tet^r)]$ | Agilent Technologies |
| *E. coli* S17-1 | $Tp^R Sm^R recA thi pro hsdR^- M^+ RP4 2-Tc::Mu-Km::Tn7 \lambda pir$ | [39] |
| *E. coli* NEB5α | $fhuA2 \Delta(argF-lacZ)U169 phoA glnV44 \Phi80 \Delta(lacZ)M15 gyrA96 recA1 relA1 endA1 thi-1 hsdR17$ | New England Biolabs |
| PAO1 | *P. aeruginosa* PAO1 (Iglewski strain) | Carrie Harwood |
| PAO1 *cheA-mTq* | *cheA-mTq* at the native chromosomal site in PAO1 | This study |
| PAO1 $\Delta cheA$ | In-frame deletion of PA1458 (*cheA*) in PAO1 | This study |
| PAO1 $\Delta cheW$ | (+9) deletion of PA1464 (*cheW*) in PAO1 | This study |
| PAO1 $\Delta che I$ | In-frame deletions of PA1456 (*cheY*), PA1457 (*cheZ*), PA1458 (*cheA*), PA1459 (*cheB*), and PA1464 (*cheW*) in PAO1 | Carrie Harwood |
| PAO1 $\Delta dipA$ | (+9) deletion of PA5017 (*dipA*) in PAO1 | This study |
| PAO1 *fliC::tn* | Transposon (lacZhah) in PA1092 (*fliC*) in PAO1. Inserted at base 820 of 1467 | University of Washington PAO1 transposon mutant collection |
| PAO1 $\Delta parC$ | In-Frame deletion of PA1462 (*parC*) in PAO1 | This study |
| PAO1 $\Delta parP$ | (+9) deletion of PA1463 (*parP*) in PAO1 | This study |

**Table S3:**
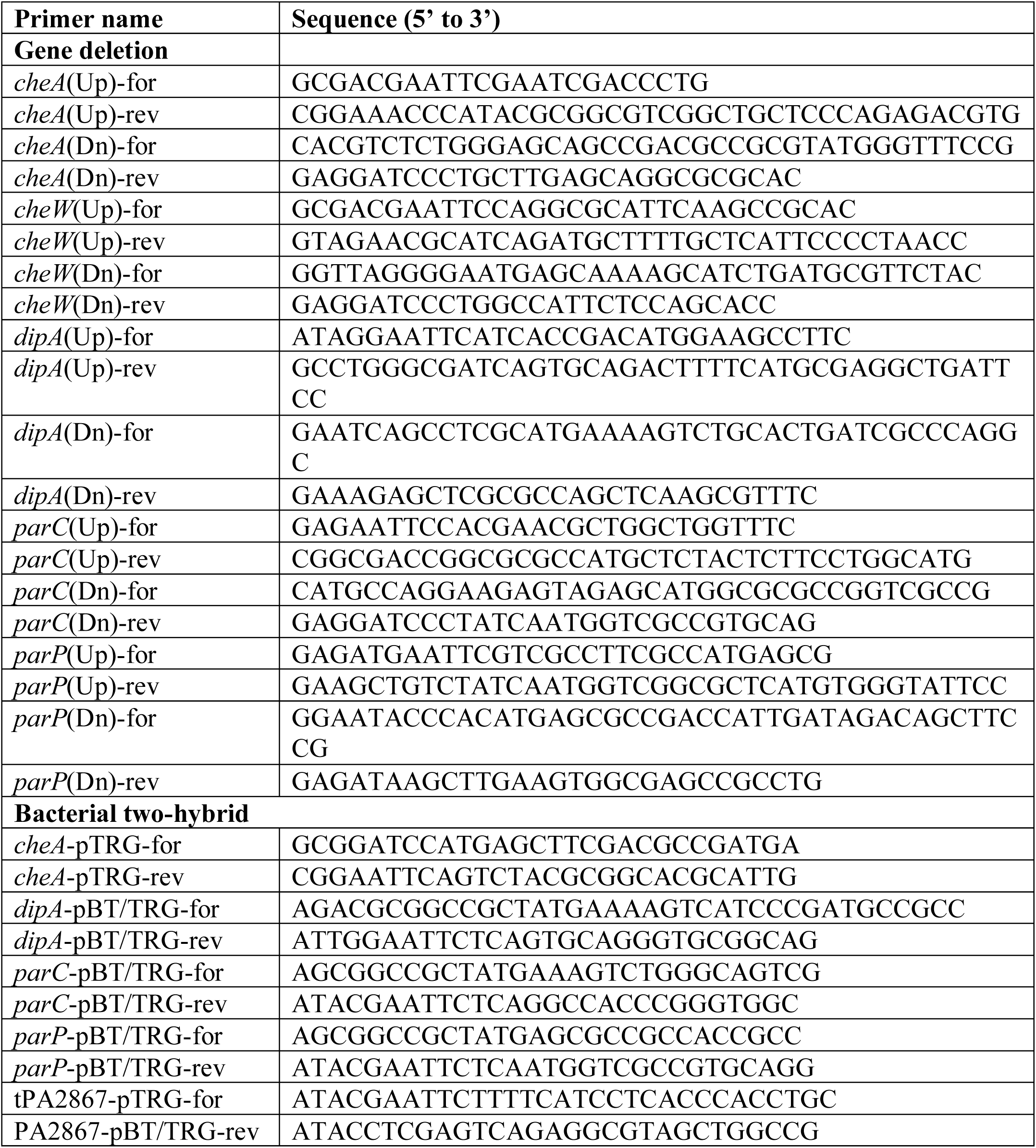
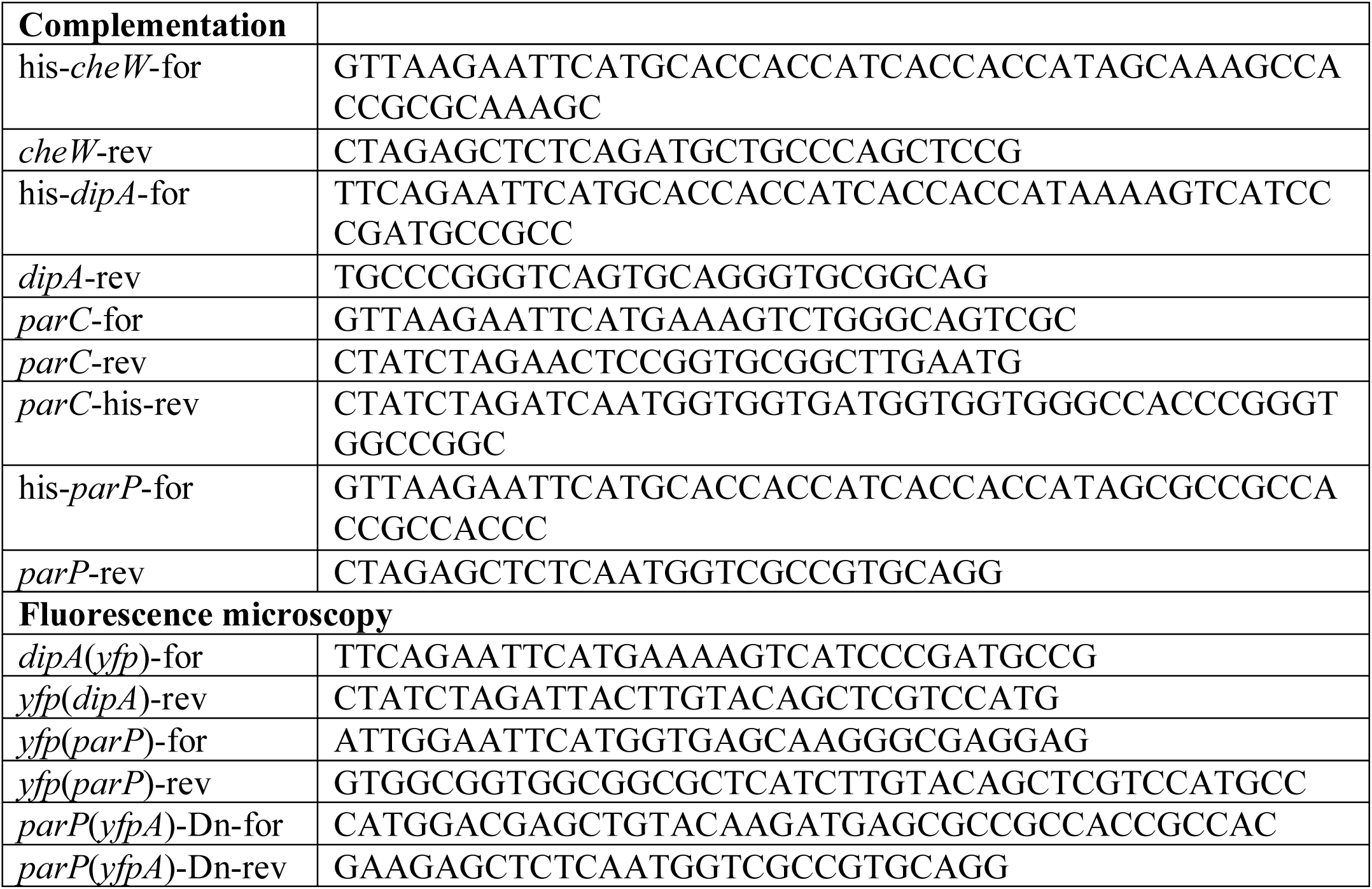
Primers used in this study

